# A Random Forest Classifier for Protein-Protein Docking Models

**DOI:** 10.1101/2021.06.23.449420

**Authors:** Didier Barradas-Bautista, Zhen Cao, Anna Vangone, Romina Oliva, Luigi Cavallo

**Affiliations:** Kaust Catalysis Center, Physical Sciences and Engineering Division, King Abdullah University of Science and Technology (KAUST), Thuwal 23955-6900, Saudi Arabia; Pharma Research and Early Development, Therapeutic Modalities, Roche Innovation Center Munich, Large Molecule Research, Nonnenwald 2, 82377 Penzberg, Germany; Department of Sciences and Technologies, University Parthenope of Naples, Centro Direzionale Isola C4, I-80143 Naples, Italy

## Abstract

Herein, we present the results of a machine learning approach we developed to single out correct 3D docking models of protein-protein complexes obtained by popular docking software. To this aim, we generated a set of ≈7×10^6^ docking models with three different docking programs (HADDOCK, FTDock and ZDOCK) for the 230 complexes in the protein-protein interaction benchmark, version 5 (BM5). Three different machine-learning approaches (Random Forest, Supported Vector Machine and Perceptron) were used to train classifiers with 158 different scoring functions (features). The Random Forest algorithm outperformed the other two algorithms and was selected for further optimization. Using a features selection algorithm, and optimizing the random forest hyperparameters, allowed us to train and validate a random forest classifier, named CoDES (COnservation Driven Expert System). Testing of CoDES on independent datasets, as well as results of its comparative performance with machine-learning methods recently developed in the field for the scoring of docking decoys, confirm its state-of-the-art ability to discriminate correct from incorrect decoys both in terms of global parameters and in terms of decoys ranked at the top positions.

## 1 Introduction

Protein-protein interactions (PPIs) are pivotal to the most diverse biological processes, including signal transduction, electron transfer and immune response. Proteins interact with each other in a specific manner and increasing evidence is revealing that perturbation of such interactions frequently leads to defective phenotypes. Mutations associated to human genetic disorders, for instance, commonly alter the interaction between proteins, rather than their folding and stability (Sahni *et al.*, 2015). *Moreover, it has been recently shown for thirty-three cancer types that amino acid substitutions in interacting proteins are significantly more frequently found in interface regions rather than in non-interface ones* (Cheng *et al.*, 2021). Targeting PPIs has thus become an essential strategy for the development of new drugs (Lu *et al.*, 2020). For fully understanding the functional and dysfunctional biological processes driven by PPIs and to possibly interfere with them, the knowledge at atomic detail of the involved protein-protein complexes would be required. However, a dramatic disproportion still exists between the number of experimental structures solved for protein complexes and the number of characterized PPIs (Mosca *et al.*, 2013). *It has been estimated that only 7% of the known human interactome is structurally characterized to date.* In this scenario, the structure prediction of protein-protein complexes by molecular docking becomes the method of choice in numerous cases of interest (Lensink *et al.*, 2018) (Harmalkar and Gray, 2021).

In a docking process, the searching step, consisting in the generation of 10^3^–10^5^ alternative poses to sample the conformational landscape, is followed by the scoring step, where the generated poses are evaluated in order to single out the correct solutions within the pool of generated poses. As the energetics governing the interaction is highly complex, scoring is a critical step in docking and has in fact become object of blind assessment in a separate challenge of the CAPRI (Critical Assessment of PRedicted Interactions) experiment (Lensink et al. 2007).

Traditionally, scoring functions for protein-protein docking poses are either energy-based or knowledge-based; the former approach using a linear combination of energy terms (which can be physics-based and/or empirical), the latter one embodying in atom-atom or residue-residue potentials the statistical occurrences observed in databases of experimental 3D structures (Huang, 2014; Moal *et al.*, 2013; Vangone *et al.*, 2017). However, over the years, a wide variety of biophysical functions and potentials have been developed, some of them combining the above potentials into a hybrid approach (Pierce and Weng 2007; Andrusier et al. 2007; Vreven et al. 2011) or integrating them with evolutionary information (Andreani *et al.*, 2013), some others based on alternative approaches, such as the consensus of the inter-residue contacts at the interface of the complex (Chermak *et al.*, 2016; Oliva *et al.*, 2013).

Nowadays, over 100 scoring functions are available from the CCharPPI web server (Moal *et al.*, 2015), while more potentials can be obtained from other public sources. These are all descriptors of the protein-protein complexes, which can be in principle combined to gain an improved performance in assessing the quality of predicted 3D models. Some of them have indeed been successfully used/combined in machine learning (ML) models recently proposed for evaluating protein-protein docking models (DMs) (Cao and Shen, 2020; Cao and Shen, 2020; Geng *et al.*, 2020; Moal *et al.*, 2017; Wang *et al.*, 2020).

Herein, we present the results of a ML approach we developed to exploit all the scoring functions that we could collect from public sources (Figure 1). To this aim, we generated a set of ≈7×10^6^ DMs with three different docking programs for the 230 complexes in the protein-protein interaction benchmark 5 (BM5), which have high-quality structures available along with the unbound structures of their components (Vreven *et al.*, 2015). Three different ML approaches were used to train classifiers with 158 different scoring functions (features). The large number of decoys used for training and validation, together with the high number of different features to describe them, is unprecedented in the field. After validation, features selection and hyperparameters optimization, a random forest classifier was selected, exhibiting the most satisfactory performance, that we named CoDES (COnservation Driven Expert System). Testing of CoDES on independent datasets, as well as results of its comparative performance with ML methods recently developed for the scoring of docking decoys in the field, confirm its state-of-the-art ability to discriminate correct from incorrect decoys both in terms of global parameters and in terms of decoys ranked at the top positions.

**Figure 1.**
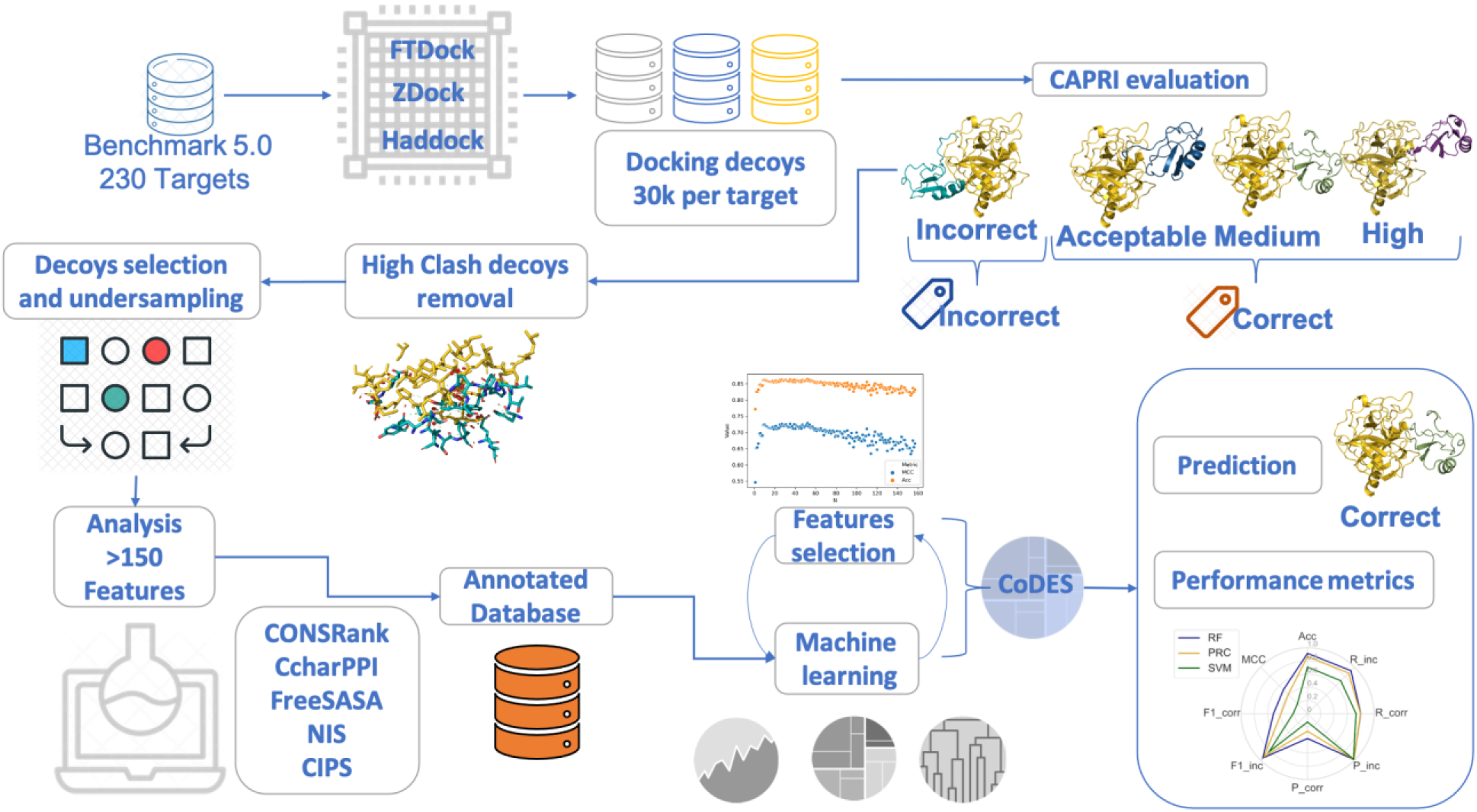
An overview of the classifier development. ≈10^7^ docking decoys were generated for the BM5 targets with FTDock, ZDOCK and HADDOCK. Based on the CAPRI quality criteria, they were labeled as correct or incorrect. Decoys with a high number of clashes were removed. By undersampling, balanced and unbalanced (in terms of included correct and incorrect decoys) datasets were derived for training and validation. For each decoy, 158 features (scoring functions and complex descriptors) were calculated. Three different machine-learning algorithms were applied, from which, upon validation, the Random Forest (RF) one was selected. The optimized RF classifier, CoDES, was then independently tested.

## 2 Materials and Methods

### 2.1 Machine learning approaches and evaluation metrics

We performed a data analysis and employed ML algorithms within the scikit learn python library (Pedregosa *et al.*, 2011; Varoquaux *et al.*, 2015) and within pyplot/seaborn for visualization (Hunter, 2007; Waskom *et al.*, 2018). We selected three classic ML algorithms, Random Forest (RF), Support Vector Machines (SVMs), and single-layer Perceptron (PRC). The RF consists in a set of decision trees, SVMs are hyperplanes that try to best separate the feature space, and PRC is a one-layer neural network. We initially trained these classifiers with default parameters.

We calculated global standard metrics to evaluate the ML predictions in our test sets as follows:

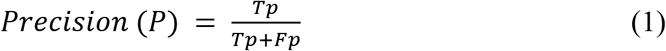

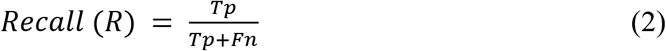

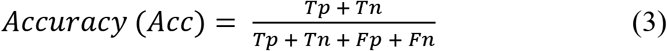

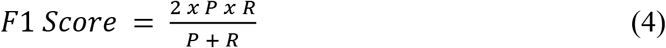

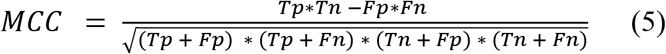

where Tp stands for true positives, Tn for True negatives, Fp for False positives, Fn for False negatives. For the independent testing on the Score_set and the IRaPPA benchmarks (see below), we also used the success rate, that is the number or percentage of correct solutions within the top *N* positions.

### 2.2 Docking models and datasets

#### 2.2.1 Docking models (DMs) generation and quality assessment

For each of the 230 protein-protein complexes (targets) in the protein-protein docking Benchmark 5 (BM5) (Vreven *et al.*, 2015), we generated a total of 30,000 DMs with FTDock (Gabb *et al.*, 1997), ZDock (Chen *et al.*, 2003) and HADDOCK (de Vries *et al.*, 2007; Dominguez *et al.*, 2003). To produce the DMs, we ran these rigid-body docking programs with the unbound structures according to the following setup. FTDock was used to generate a total of 10,000 DMs per target with electrostatic interactions switched on, a grid cell size of 0.7 Å, and surface thickness of 1.3 Å. ZDOCK, 3.0.1 was used to generate a total of 54,000 DMs per targets, of which we kept the top-scoring 10,000 ones. Finally, HADDOCK was used to generate 10,000 DMs per target by applying the HADDOCK rigid body step default parameters.

The quality of the generated DMs was assessed following the CAPRI (Critical Assessment of PRedicted Interactions) protocol (Mendez *et al.*, 2003). To this aim, for each DM we calculated three parameters: f(nat), which represents the fraction of contacts in the target (the native structure experimentally determined) that is reproduced in the DM, where a contact is defined as any pair of atoms from the ligand (smaller size protein) and the receptor (larger size protein) within 5 Å to each other; L-rms, which is the root mean square deviation (RMSD) of the backbone atoms of the ligand after optimally superimposition of the receptor in the DM and the target structure; and I-rms, i.e. the RMSD of the backbone atoms of all interface residues after they have been optimally superimposed, where interface residues are those having at least an heavy atom within 10 Å of any atom of the binding partner. Based on the values of the above parameters, DMs were classified, in order of increasing quality, as Incorrect, Acceptable, Medium- and High-quality, as reported in Table S1. All the DMs are available at (https://doi.org/10.5281/zenodo.4012018).

An initial screening of DMs was performed by removing the 17 targets for which no correct DM was identified in the 30,000 generated ones. For the 213 remaining targets, we classified overall 246 high quality, 7,146 medium quality, 42,815 acceptable and 6,339,793 incorrect DMs. For these targets we calculated the number of clashes for each of the 30,000 generated DMs. As in CAPRI, clashes were defined as contacts between non-hydrogen atoms separated by a distance below 3 Å. For each target, we discarded DMs with a number of clashes greater than the average number of clashes plus two standard deviations for that target. Overall we discarded 243,061 DMs, which left us with a total of 6,146,939 DMs, on average 28,859 ± 218 DMs per target.

#### 2.2.2 Balanced dataset for ML analysis

Since databases containing DMs generated by docking software are highly unbalanced, to train the ML algorithms we built a “balanced” dataset by applying a random undersampling selection of the incorrect DMs to match the number of correct ones. We performed this undersampling case by case, meaning that for each target we matched its number of correct DMs with the same number of randomly sampled incorrect DMs. In this way we produced a balanced dataset, Bal-BM5, where each target has the same number of correct and incorrect DMs. The number of DMs per target is quite variable in the Bal-BM5 dataset, as it depends on the number of correct DMs available for each specific target. The number of correct DMs per target ranges from 2 to 600, with the same number of incorrect ones. On average the number of DMs per target is 346 (with a standard deviation of 358), while the median value is 214. Only DMs that passed the check on clashes were included in the Bal-BM5 dataset.

The targets and DMs selected in the above steps were separated into training and validation sets using two different strategies. As a first strategy, all the 161 targets included in the benchmark 4 (BM4) were assigned to the training set, Bal-BM4, while the remaining 52 targets, corresponding to the BM5 update, relatively to BM4, were assigned to the validation set, Bal-BM5up. Alternatively, for the cross-validation (CV) approach, we randomly selected 64 targets from the Bal-BM5 dataset to produce a validation set, while the remaining 149 targets were used for training. This process was repeated 10 times.

#### 2.2.3 3K unbalanced dataset for ML analysis

As the typical case in docking challenges is having a small fraction of correct DMs in the total ensemble of DMs, we decided to build unbalanced datasets for validation and testing of the trained ML models. To this end, we selected 3,000 DMs per target, out of those generated in the docking step and that passed the check on clashes. For each target we selected up to 600 DMs with acceptable or better quality, and picked up randomly incorrect DMs until 3,000 DMs in total were collected into the 3K-BM5 dataset. We fixed a maximum value for the number of correct DMs, 600, to avoid biasing the training of the ML algorithms towards a few cases with a too large number of correct DMs. Strategies similar to those used to split the whole Bal-BM5 dataset into training and validation datasets were used to extract validation datasets from the 3K-BM5 dataset. Specifically, the 52 targets corresponding to the BM5 update were assigned to the 3K-BM5up dataset or, as an alternative, for the cross-validation approach 64 targets were randomly selected to produce a validation dataset, repeating this process ten times.

#### 2.2.4 Independent CAPRI Score_set and IRaPPA test sets

For an independent testing, we used the CAPRI Score_set, made of 15 targets from the CAPRI Rounds 13-26, with a total of 19,013 DMs, of which 2,166 (11%) are correct. This benchmark set contains DMs predicted by 47 different predictor groups including web servers, which use different docking and scoring procedures, being thus highly diverse, and include dimers and multimers (Lensink *et al.*, 2016). For 13 of these targets at least one correct solution is available. As another independent test dataset, we used the DMs generated with ZDOCK by the IRaPPA authors for validating their method,(Moal *et al.*, 2017) for all the 55 targets (30 of them including no correct solution) in the BM5 update. They include a total of 25,728 decoys (468 on average per target), of which 1,097 (4.3%) classified as correct.

### 2.3 Scoring functions (features)

We analyzed DMs of the 213 targets from BM5 with a total of 157 descriptors (features). Ninety-two of such features come from the CCharPPI server (Moal *et al.*, 2015) and mostly consist in physics-based or empirical energy terms (see Table S2). We did not consider 16 CCharPPI scoring functions that failed to evaluate more than 30% of the total number of DMs. Of the used 92 CCharPPI functions, 37 failed on average on 4% of the analyzed DMs. In these cases, for the DMs missing the corresponding score, the mean value of that scoring function for the corresponding target was assumed. Of the 65 non CCharPPI features we considered, 32 are calculated by our tools CONSRANK (Oliva *et al.*, 2015; Oliva *et al.*, 2013) and COCOMAPS (Vangone *et al.*, 2012; Vangone *et al.*, 2011), and consist of the consensus CONSRANK score, based on the frequency of inter-residue contacts at the interface, and of the number of inter-residue contacts per class of involved amino acids (apolar, polar, aliphatic, aromatic, charged), using a 5 Å cut-off (see Table S2). Twenty-eight more features come from CIPS (combined interface propensity per decoy scoring (Nadalin and Carbone, 2018)), and represent sums and averages of the CIPS score over the different classes of residues at the interface (polar, apolar, polar aliphatic, aromatic and charged); 3 from the buried surface area (BSA), total, polar and apolar, calculated with FreeSASA (Mitternacht, 2016), and 2 are non-interacting surface (NIS) terms (Kastritis *et al.*, 2014), polar and apolar, calculated by Prodigy (Vangone and Bonvin, 2015). The above features were used to train the ML algorithms. The values of theses scoring functions were normalized as Z-scores using Eq. 6:

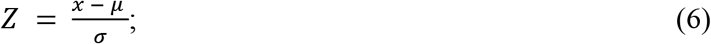

where *μ* and *σ* are the mean and the standard deviation of the corresponding scoring function on the training set.

### 2.4 Feature selection and optimization of the random forest classifier

It is widely acknowledged that a feature selection step, i.e. the removal of irrelevant and weakly relevant features, may improve the performance of a predictor, in terms of accuracy, simplicity and/or speed (John *et al.*, 1994). For the feature selection, we used the sequential forward feature selection method (Blum and Langley, 1997; Kudo and Sklansky, 2000; Marcano-Cedeno *et al.*, 2010) (Schenk *et al.*, 2009). Specifically, we followed a bottom-up search procedure gradually adding features selected on the basis of their importance in the RF classifier. Accuracy and the Matthews correlation coefficient (MMC) were used as prediction evaluation metrics.

To fine tune the RF algorithm we performed a series of tests to find the best combination of hyperparameters. Specifically, we varied over a regular grid the number of trees, the minimum number of samples to split the node, and the weight assigned to correct and incorrect models, as these parameters of the RF algorithm have a clear influence on the final prediction. To reduce the number of computational tests, the three hyperparameters were optimized independently.

## 3 Results and Discussion

### 3.1 Training and optimization of the ML classifiers

#### 3.1.1 Selecting a ML algorithm

Three ML algorithms, Random Forest (RF), Support Vector Machine (SVM) and Perceptron (PRC), were initially used to train a classifier for protein-protein DMs, using DMs for 213 targets from the protein-protein interaction benchmark (BM) (Hwang *et al.*, 2010) (Vreven *et al.*, 2015)). This benchmark (Hwang *et al.*, 2010) (Vreven *et al.*, 2015)) is the gold standard in the field, widely used for testing protein–protein docking and scoring methods. It consists of non-redundant, high-quality structures of enzymes containing, antibody-antigen and other types of protein–protein complexes. Of the 213 complexes considered by us, 52 were added in the latest version 5 (Vreven *et al.*, 2015) and are referred to in the following as BM5-update. All the other 161 complexes were already included in the benchmark version 4 and are referred to in the following as BM4.

Two main DMs datasets were used for them, the former (the balanced dataset) presenting a variable number of DMs per target (346 on average) but equally distributed between correct and incorrect DMs, the latter (the 3K dataset) consisting in a dataset of 3,000 DMs per target, with a variable number of correct and incorrect DMs (see Methods). As a consequence of their definition, the 3K dataset is ≈9 times larger than the balanced one and presents a number of incorrect DMs ≈15-fold higher. All the DMs in these datasets were generated by us with three different docking programs: FTDock, ZDock and HADDOCK (see Methods), and include at least one correct solution per target.

The default training of the classifiers was performed on the targets of the BM4 balanced dataset, Bal-BM4, while validation was performed on the targets included in the BM5 update both using the 3K and the balanced datasets, 3K-BM5up and Bal-BM5up. The comparative performance of the three classifiers trained on Bal-BM4 and tested on the Bal-BM5up and 3K-BM5up datasets is reported in Figures 2 as radar plots. The RF classifier clearly outperforms the other two classifiers on both validation datasets. The difference in performance is particularly apparent on the 3K-BM5up dataset, especially in terms of precision at retrieving the correct solutions, recall of the incorrect solutions and overall accuracy, all reflected in the significantly higher MCC of the RF classifier (all the values are reported in Tables S3). As the 3K-BM5up set is closer than the Bal-BM5up to a real-case scenario, the performance of the classifier on it deserves special attention. We observe that the lowest metric for it is in the precision in classifying correct DMs, implying a high number of false positives. This low precision clearly impacts the corresponding F1 score (F1_corr).

**Figure 2.**
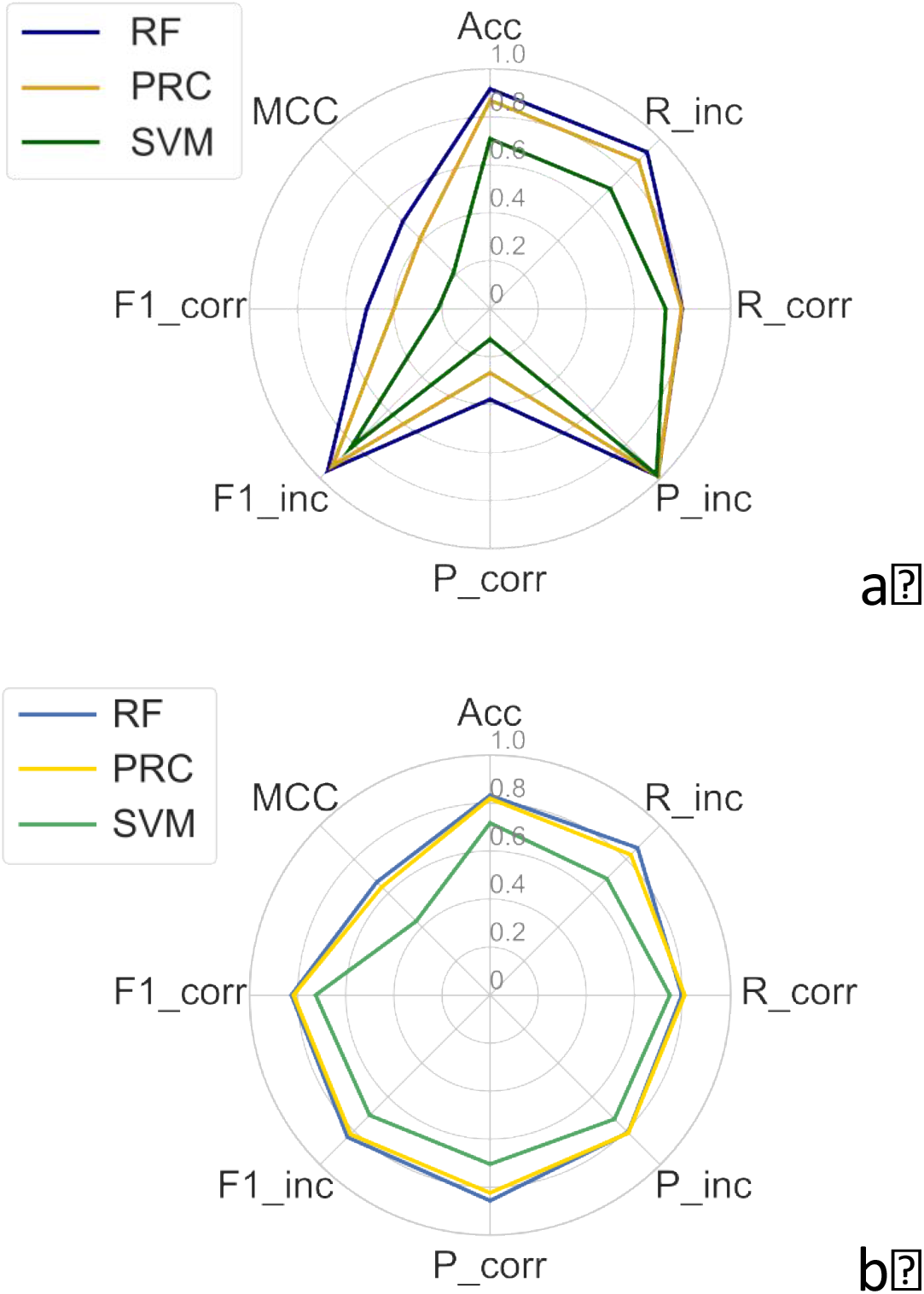
Radar plot of the performance metrics for the RF, SVM and PRC classifiers on the 3K-BM5up (**a**) and Bal-BM5updataset (**b**) as test sets. All the classifiers were trained with all the features available on the 161 targets in BM4 and tested on the 52 targets in the BM5 update (balanced dataset). “P” stands for precision, “R” for recall, “Acc” for accuracy and “F1” for F1-score, while MCC is the Matthews’ correlation coefficient. “Corr” abbreviates “correct” and “inc” abbreviates “incorrect”.

The performance of all the classifiers on the Bal-BM5up dataset is quite satisfactory, with MCC values ranging between 0.44 and 0.67, and differences between them become less dramatic. However RF still significantly outperforms the other classifiers, especially the SVM one.

#### 3.1.2 Cross-validation testing on the random forest classifier

As the RF classifier was more promising than the SVM and PRC ones, we further tested it using a cross-validation (CV) approach. The results of a 10-fold CV on the whole Bal-BM5 dataset (Figure 3) are compared with those previously obtained, by training on the Bal-BM4 dataset and testing on the 3K-BM5up and Bal-BM5up datasets. Performance obtained with the two validation approaches are quite similar, both on the balanced and on the 3K dataset. For the balanced set, the larger differences are in the recall, which is decreased (from 0.797 to 0.720) for correct and increased (from 0.867 to 0.925) for the incorrect solutions. For the 3K set, the cross-validation approach only affects the metrics for retrieving the correct solutions, by slightly increasing the precision (from 0.378 to 0.504) while decreasing the recall (from 0.798 to 0.734), eventually resulting in slightly enhanced F1 score and MCC values (to 0.802 and 0.584, respectively).

**Figure 3.**
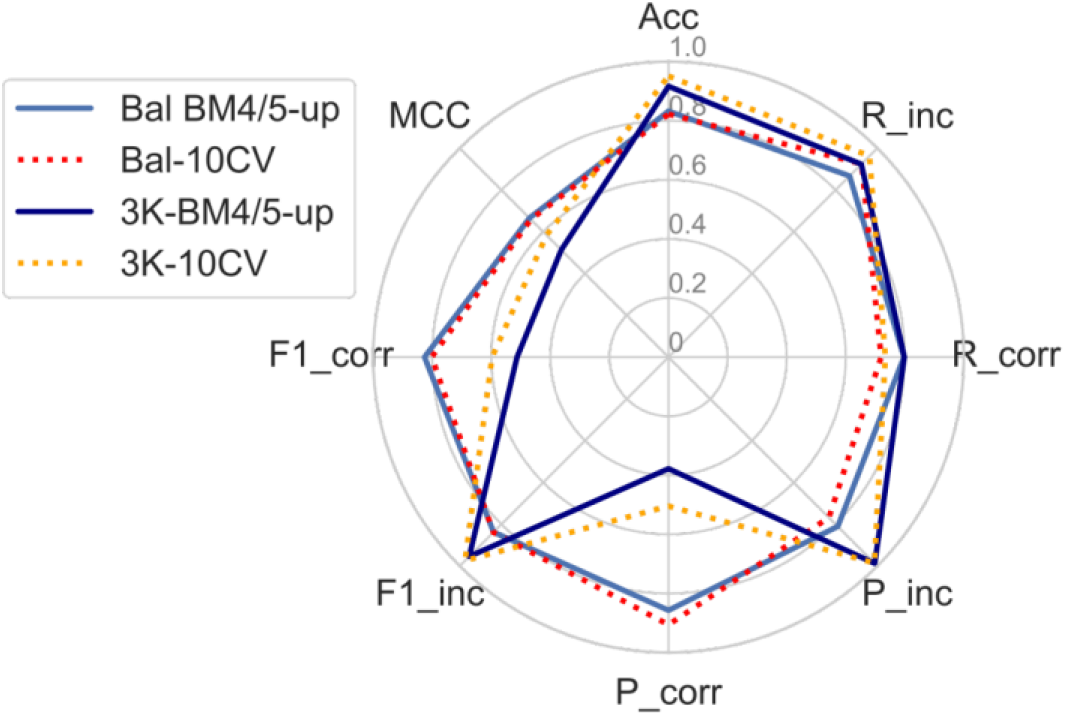
Radar plot of the performance metrics for the RF when trained with a 10-fold cross-validation approach on the balanced set and tested on the balanced and 3K sets (dotted lines). For the sake of comparison, the RF performance according to the BM4/BM5-update approach is also reported (the solid dark and light blue lines are the same shown in Figure 2).

#### 3.1.3 Selection of features for the RF classifier

An advantage of the RF is the possibility of retrieving the features important for classification. We used a forward selection protocol to determine the threshold of importance for a feature to influence the classification. We evaluated the importance of the features for the two different training approaches described before, which is training on Bal-BM4 with validation on the Bal-BM5up and 10-fold CV on Bal-BM5. Monitoring performance metrics showed that 1% is a suitable importance threshold to optimize the classifier performance (Figure 4).

**Figure 4.**
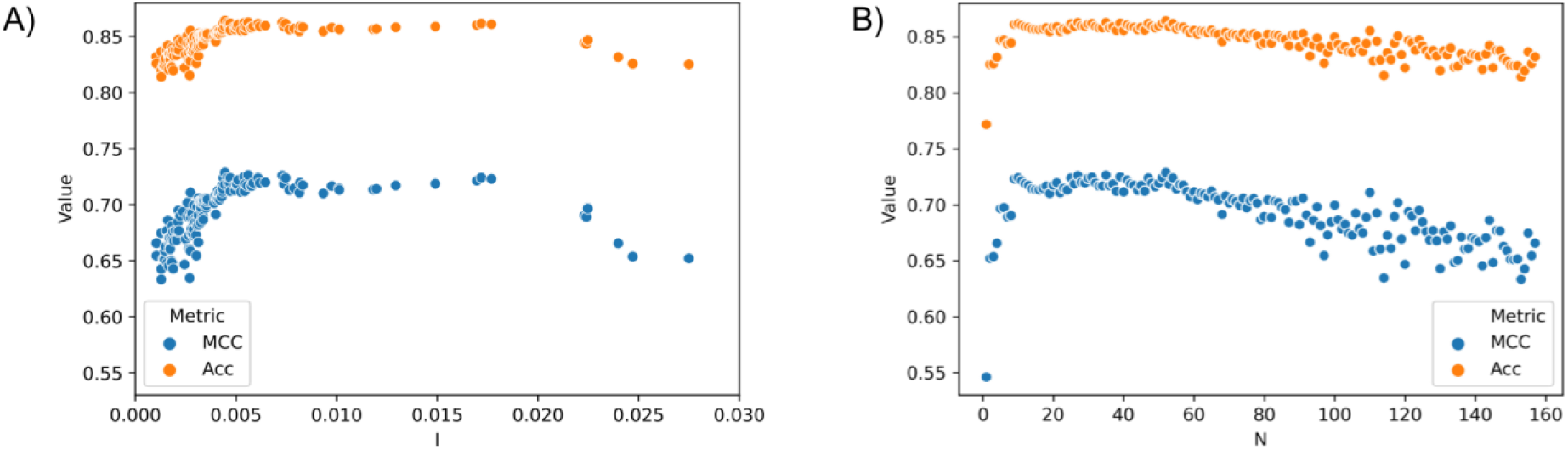
Scatter plot of the accuracy (Acc) and the Matthews’ correlation coefficient (MCC) for the RF classifier trained on the Bal-BM4 and tested on the Bal-BM5up for the forward feature selection. “I” stands for the features importance, while “N” is the number of features added. **A**) The plot shows the change for the two metrics upon adding features with importance ranging from 0 to 0.03. Both accuracy (Acc) and the Matthews correlation coefficient reach a plateau in the performance after a importance of 0.01. **B**) The plot shows the change for the two metrics when adding sequentially an increasing number (N) of different features from the highest to the lowest importance. After adding 10 to 20 features (corresponding to an importance of ≈0.01) the two metrics are maximized, reach a plateau and then, after ≈50 features are added, slowly decrease.

Overall, 16 features with an importance above 0.01 were common to the two selection approaches (see Table S4) and were selected for further classification training. The ranking of the top ten important features was virtually the same for the two approaches, as can be seen in Table 1.

**Table 1.**
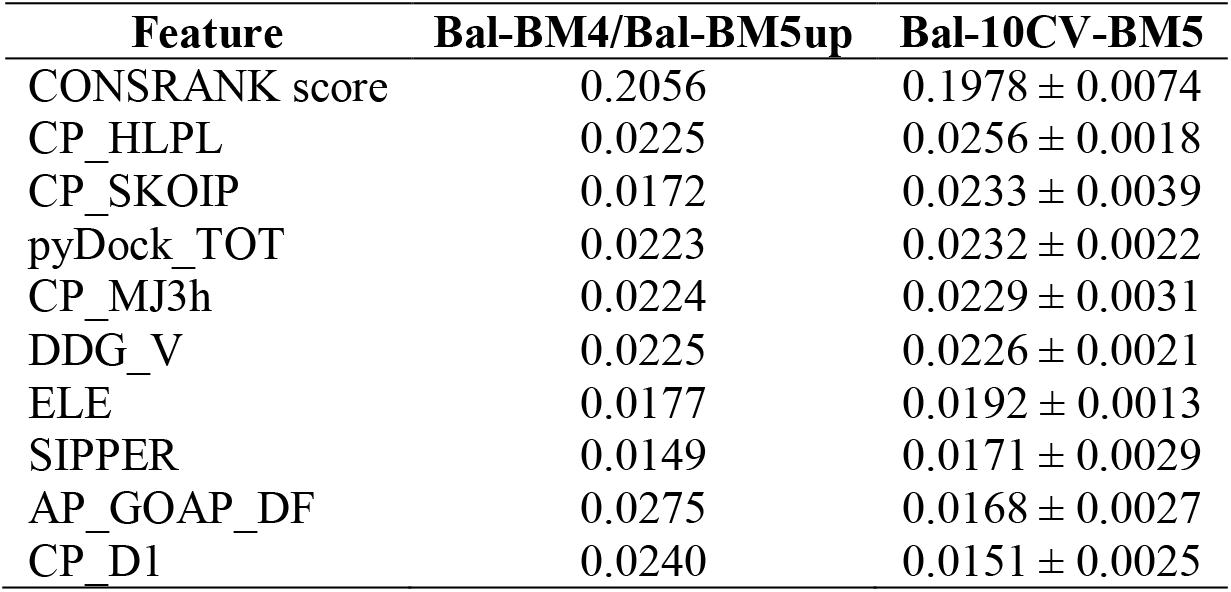
The top ten features with highest importance are reported for the RF classifier trained on the balanced sets with a BM4/BM5-update and 10-fold cross-validation approach. Features are sorted based on values of the second column. The mean importance value and associated standard deviation is reported in the last column.

One feature has an importance around 0.2 for both the approaches, being one order of magnitude larger than any other one. It is the CONSRANK score, a consensus score, which reflects the conservation (or frequency) of the inter-residue contacts featured by a given model in the whole decoys ensemble it belongs to (Chermak *et al.*, 2015; Oliva *et al.*, 2015; Oliva *et al.*, 2013), widely tested on CAPRI targets (Barradas-Bautista *et al.*, 2020; Lensink *et al.*, 2019; Lensink *et al.*, 2016; Vangone *et al.*, 2013). Among the top ten scoring functions, after CONSRANK, we find CP_HLPL (Garcia-Garcia *et al.*, 2010; Pokarowski *et al.*, 2005), CP_SKOPI (Lu *et al.*, 2003), pyDock_TOT, CP_MJ3h, DDG_V, ELE (electrostatics), SIPPER, AP_GOAP_DF and CP_DI, all obtained from the CCharPPI server (Moal *et al.*, 2015). They include 3 residue contact/step potentials (Garcia-Garcia *et al.*, 2010; Pokarowski *et al.*, 2005) (Lu *et al.*, 2003), 1 residue distance-dependent potential (Liu and Vakser, 2011), 1 atomic distance-dependent potential (in particular the DFIRE term in the GOAP energy (Zhou and Skolnick, 2011), 2 composite scoring functions (Cheng *et al.*, 2007; Grosdidier and Fernández-Recio, 2008; Pons *et al.*, 2011), the electrostatic component of the PyDock scoring function, and a miscellaneous scoring function, consisting in a microscopic surface energy model derived from mutation data (Moal *et al.*, 2015).

Selected features thus include statistical potentials, energy-based potentials as well as consensus scores, thus including scoring functions from all the representative approaches in the field (Huang, 2014; Moal *et al.*, 2013; Moal *et al.*, 2013; Vangone *et al.*, 2017).

#### 3.1.4 Tuning of the random forest classifier hyperparameters

The hyperparameters that can be optimized to improve the performance of a RF classifier are: the number of trees, the minimum number of samples to split the nodes and the weights assigned to each class. We tested different values for all these three hyperparameters, finding out that the number of trees does not affect significantly the performance of our classifier; therefore it was kept at the default value of 100 in sklearn. The optimal value for the minimum number of samples to split the nodes was instead found to be 10, versus a default value of 2 from sklearn, while the optimal weights for the class 0 (incorrect) and class 1 (correct) were identified as 1.2 and 0.1, respectively. For details see the Supplementary Materials. These optimal values were used for the training of the final classifier (see below).

#### 3.1.5 Training and testing of the final classifier (CoDES)

After selecting the features and optimizing the RF hyperparameters, we trained our final RF classifier with selected features and optimal hyperparameter values on the Bal-BM4 dataset. We named this classifier CoDES (Conservation Driven Expert System) and tested it in different conditions. First, we evaluated the performance of CoDES using the unbalanced validation set 3K-BM5up, showing that it outperforms the RF classifier trained on all the features and with default parameters, as concerns the recall and precision in retrieving the correct solution (increased respectively from 0.80 to 0.90 and from 0.38 to 0.46, see Figure 5a), which are then reflected in the MCC, which is increased from 0.51 to 0.61.

**Figure 5.**
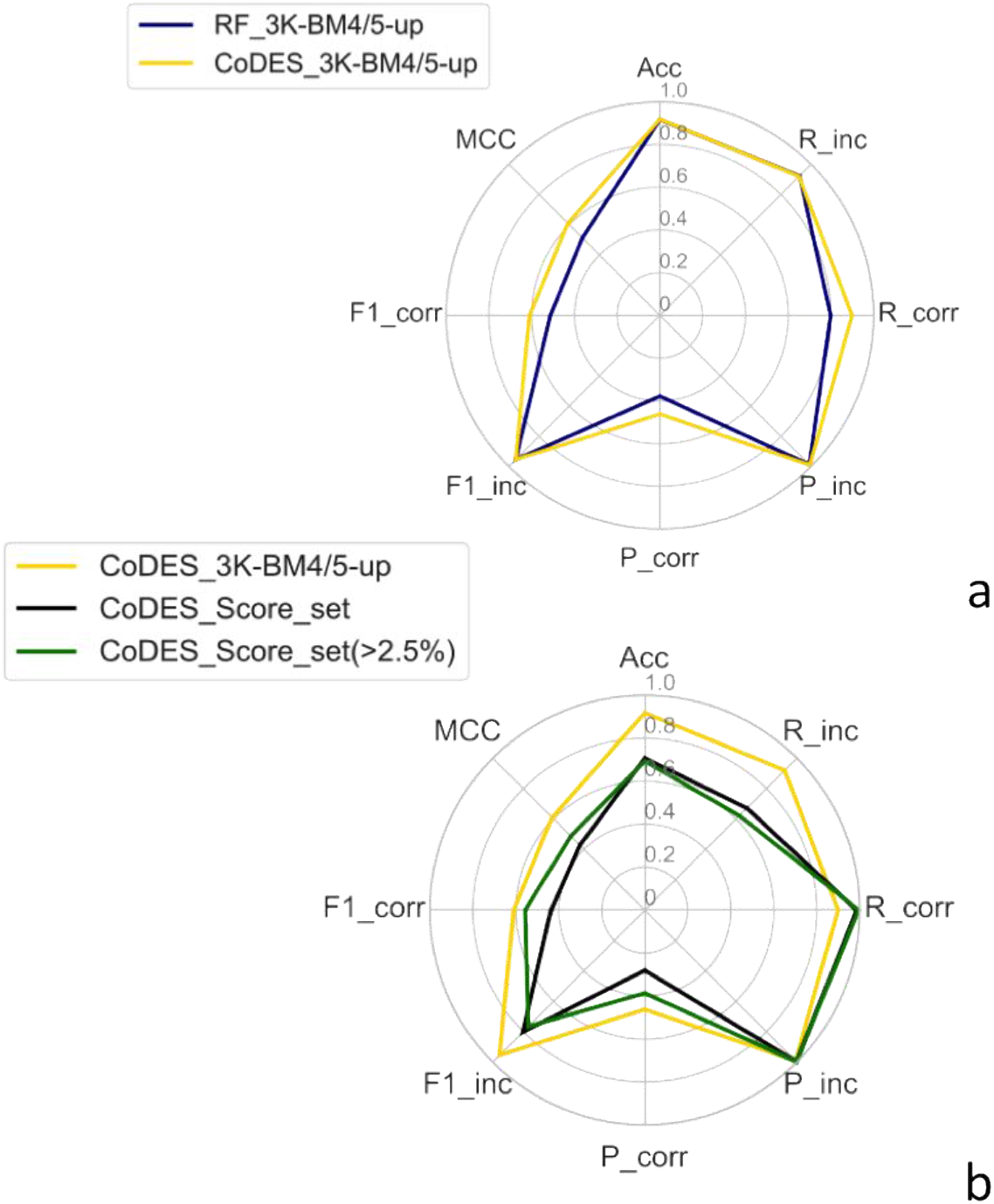
Performance of CoDES on unbalanced datasets. **a)** Radar plot of the performance metrics for CODES on the unbalanced validation set (BM5 update, golden line). For the sake of comparison, the original RF classifier performance according to the BM4/BM5-update approach is also reported (the solid dark line is the same shown in Figures 2–3). **b)** Radar plots of the performance metrics for CoDES on the complete unbalanced Score_set (black line) and on the Score_set targets with over 2.5% of correct models (green line).

In addition, for an independent testing we used Score_set, a set of models for 15 targets, submitted by several predictors to various rounds of the CAPRI experiment, with an average of 1267 models per target. As expected, the performance metrics on Score_set are lower than on 3K-BM5up. Score_set is indeed a particular difficult test set, highly unbalanced as it includes 0 correct solutions for 2 targets (T36 and T38) and a percentage of correct models ≤ 2.5% for 5 more targets (T30, T32, T35, T39, T54). Results are shown in Figure 5b in the form of a radar plot. Again, due to the imbalance on both sets, retrieval of correct models is the real challenge, as reflected by the corresponding F1-score (0.43). While the recall of correct solutions is even enhanced for the Score_set (0.98 vs 0.90, Figure 5), the precision drops from 0.46 to 0.28. This is reflected in the overall MCC value, which however stays at a satisfying value of 0.43. When excluding from the analysis the highly unbalanced targets, i.e. the targets featuring no more than 2.5% of correct DMs, the F1-score for the correct class retrieval increases to 0.56, due to a gain in the corresponding recall, and especially of the precision, increasing to 0.39. The MCC on these targets is as high as 0.49.

#### 3.1.6 Comparative performance of CoDES with other ML methods

A widely-used metric for evaluating the performance of a scoring function is the ability to have correct DMs ranked at the top-*N* positions for each target. This kind of metric responds to the demand from researchers to have a limited number of putative solutions to be further explored by computational or experimental approaches. It can also been referred to as the “success rate” and is the standard metric in the CAPRI experiment. Several recently developed ML methods for the scoring of DMs have been evaluated/*tested* in terms of success rate, therefore we will use in the following such a metric for comparing the CoDES performance with them, on datasets for which their results have been made available.

Performance in terms of top-10 success rate is available for two recently developed ML algorithms (Geng *et al.*, 2020) (see Table 2), for the 13 targets in the Score_set benchmark having at least one correct model available. One algorithm, GraphRank, is a scoring function using a graph representation of the complex interface together with a measure of evolutionary conservation; the second algorithm, iScore, combines GraphRank with classical HADDOCK energy terms. For the iScore’s evalutation, the authors obtained clusters of at least 4 members, based on the fraction of common contacts (Rodrigues *et al.*, 2012), then selected the top 2 DMs of the top 5 clusters for each target. For our evaluation, instead, for each target we ranked all the DMs with CoDES, then we straightforwardly selected the 10 top ranked DMs. The same results are obviously also available for all the scoring groups participating in the corresponding CAPRI rounds.

**Table 2.** Comparison of CoDES with other machine learning classifiers, GraphRank, iScore, and with the CAPRI best performing group per target on the CAPRI score set. Only the top 10 models are evaluated. The scoring performance for each target is reported as the number of acceptable or better models (hits), followed by the number of high (indicated with ***) or medium quality models (**). In the last row, the overall performance of each method on all 13 targets is reported in a similar way. The total number of targets for which at least 1 correct/high(***)/medium(**) quality model is ranked among the top 10 is given. Note that the ‘CAPRI best’ column corresponds to results from 37 different groups. The last column reports the percentage of correct models available in the ensemble of models per target.

| Target | CoDES | GraphRank | iScore | CAPRI best | % Correct |
| --- | --- | --- | --- | --- | --- |
| **T29** | 10/7** | 4 | 4 | 9/5** | 8.0 |
| **T30** | 0 | 0 | 0 | 0 | 0.15 |
| **T32** | 1/1** | 4/1** | 4/1** | 2 | 2.5 |
| **T35** | 0 | 0 | 0 | 1 | 0.60 |
| **T37** | 10/1***/6** | 2/1** | 4/2** | 6/1*** | 6.6 |
| **T39** | 0 | 0 | 0 | 0 | 0.29 |
| **T40** | 7/3***/3** | 4/3** | 4/1*** | 10/10*** | 27 |
| **T41** | 5/2** | 8 | 10/2** | 10/2*** | 31 |
| **T46** | 2 | 3 | 4 | 4 | 1.4 |
| **T47** | 9/3***/6** | 8/5***/3** | 10/6***/4** | 10/10*** | 58 |
| **T50** | 4/2** | 0 | 4/3** | 7/6** | 9.2 |
| **T53** | 2/1** | 5/1** | 5/1** | 8/3** | 9.3 |
| **T54** | 0 | 0 | 0 | 0 | 1.4 |
| **Total** | 9/3***/5** | 8/1***/4** | 9/2***/5** | 10/4***/3** | - |

The comparative performance of CoDES, GraphRank, iScore and the best CAPRI scorer, for each target in the respective CAPRI round, is reported in Table 2. Although CoDES is a binary classifier, assigning DMs to either the correct or to the incorrect class, we kept in the table the classical CAPRI sub-classification of correct DMs in: acceptable, medium- and high-quality, in order of increasing closeness to the real structure. Table 2 presents results of such analyses.

CoDES is able to rank at least one correct solution within the top 10 positions for all the targets, but T30, T35, T39 and T54. These targets all feature a percentage of correct solutions extremely low (between 0.15 and 1.4%) and not surprisingly all the other scoring approaches also failed on them, with the exception of T35 for which one only scorer in CAPRI could rank 1 acceptable model among the top 10 positions. Overall, with its 9 targets having at least one correct model, of which 3 of high- and 5 of medium-quality, CoDES slightly outperforms GraphRank, while equaling the performance of iScore (with one target more having high-instead of medium-quality solutions top ranked). This performance is better than that of any single scorer in the corresponding CAPRI Rounds (Geng *et al.*, 2020) and only slightly worse than that of the best CAPRI scorers per target taken all together.

A further comparison can be made with other two ML methods for which less detailed results on the Score_set, i.e. the total number of targets with at least one correct, high- or medium-quality DMs ranked within the top-10 positions, are available. These methods are: a random forest classifier being a combination of eight energy potentials (Cao and Shen, 2020); and an energy-based graph convolutional network (EGCN) (Cao and Shen, 2020). Of them, the best performance was achieved by EGCN with 7/1/4 targets having at least one correct/high/medium-quality DM. CoDES thus clearly outperformed also the above methods on this test set.

Testing on Score_set showed overall that CoDES is not only able to top rank correct solutions, but also to single out DMs of medium and high-quality. This outcome, maybe unexpected as the classifier was trained to only distinguish correct decoys from the incorrect ones independently from their detailed quality, could actually reflect overall an effective distinction between good and bad decoys, *implicit*/hidden in the features used for training the model.

A comparison similar to those above was also feasible with the recently developed IRaPPA ML method, combining physico-chemical and statistical potential descriptors for the scoring of DMs, using ranking support vector machine, an efficient approach for information retrieval (Moal *et al.*, 2017). The IRaPPA approach was developed independently for DMs from four different docking programs, including ZDOCK (Chen *et al.*, 2003). The IRaPPA performance was measured in terms of success rate on the top-1, top-10 and top-100 positions, not distinguishing between DMs of acceptable, medium- or high-quality. Therefore, we used the same metrics to compare results of CoDES with those of IRaPPA-ZDOCK and of the original ZDOCK scoring function, on the ZDOCK DMs generated for the 55 targets of the BM5-update and used by the IRaPPA authors for the method training and validation. For 30 of these 55 targets, however, no correct DM was included in the dataset, therefore a success rate of 45% (25 over 55 targets) corresponds to the maximum performance achievable. Results are reported in Figure 6 and show that at least 1 correct DM was ranked within the top-10 solutions for 17 targets (31% of the total 55 targets) by CoDES, for 16 targets (29%) by IRaPPA-ZDOCK and for 11 targets (20%) for ZDOCK. At the top-1 position, CoDES could locate correct DMs for 8 targets, which compare with the 3 of ZDOCK and the 11 of IRaPPA-ZDOCK, while at the top-100 positions CoDES had 23 targets with at least 1 correct DM, versus the 20 targets of ZDOCK and the 25 targets of IRaPPA-ZDOCK. CoDES therefore outperformed the original ZDOCK scoring function on this set for all the metrics, and performed slightly worse than IRaPPA-ZDOCK on it, as concerns the top-1 and top-100 positions, while having a slightly better performance at the top-10 positions. It is however worth noting that IRaPPA-ZDOCK is specific to ZDOCK, while CoDES is a non-specific method, trained and applicable on models from different sources.

**Figure 6.**
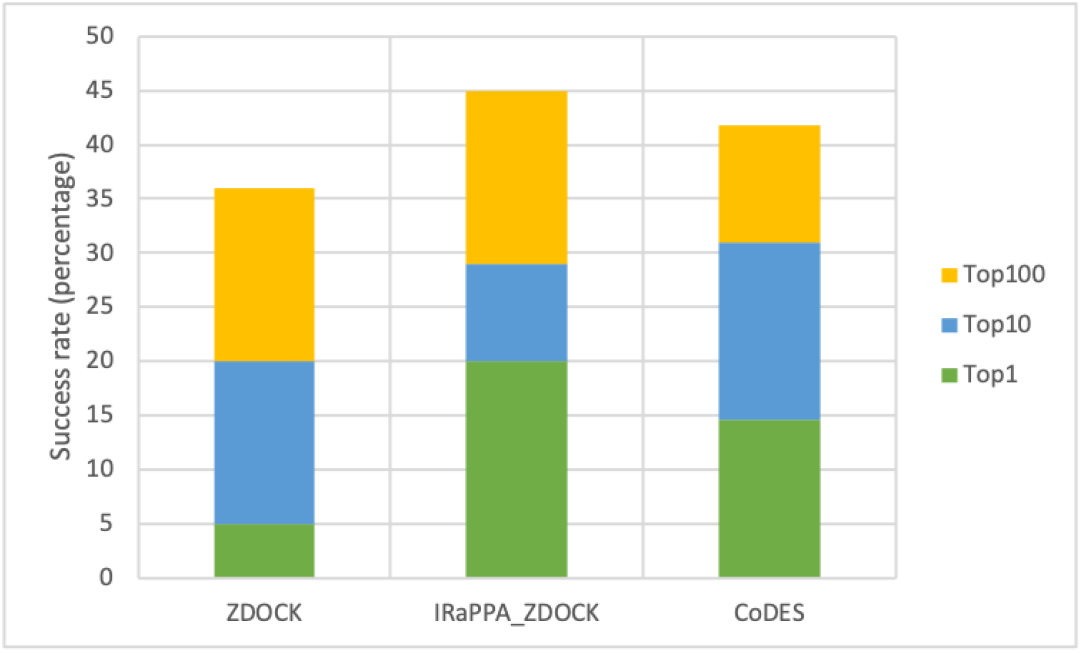
Number of BM5 targets for which at least 1 correct decoys was ranked within the top-1, top-10 and top-100 positions). The docking decoys were generated with ZDOCK by the IRaPPA authors. Only 25 of the 55 BM5 targets featured at least 1 correct decoy overall.

## 4 Conclusion

While application of ML is dramatically boosting the efficiency of structural bioinformatics tools, several ML methods are being developed for addressing one of the open challenges in the field, that is the recognition of the few correct solutions among the multitude of generated DMs for protein-protein complexes (scoring problem). Herein, we present a RF classifier selected and optimized after testing of three different ML approaches. For the initial training of the classifier, 158 features, the largest number used to the aim to date, have been collected by public sources, and an initial dataset of ≈7×10^6^ DMs has been generated with three popular docking programs, FTDOCK, HADDOCK and ZDOCK. From them, a subset of ≈7×10^4^ DMs, Bal-BM5, has been used for training and one of ≈7×10^5^ DMs, 3K-BM5, for validation. To our knowledge, these are the largest datasets used to develop a ML classifier in this field. We decided to make them open access, labeled with their respective quality assignment (incorrect, acceptable, medium- and high-quality, according to the CAPRI criteria) and complete with the values of calculated features. Properly designed as they are to represent a typical balanced or unbalanced scenario - in terms of number of correct and incorrect models -, we believe they can become reference benchmarks both for developing and comparing different scoring methods using classic empirical potentials, and for the training of ML-based methods. The final RF classifier was named CoDES (COnservation Driven Expert System), as, within the 16 selected features optimizing its performance, the one having by far the highest importance is the CONSRANK score, which represents the average conservation (frequency) of the inter-residue contacts featured by a given DM, relatively to the set of models it belongs to. Testing of CoDES on the CAPRI Score_Set showed it to outperform any single scorer in the corresponding CAPRI Rounds, and to be able to top rank not just correct but medium- and high-quality DMs. Overall testing on independent datasets resulted in CoDES equaling or exceeding the performance of the few state-of-the-art machine learning methods available in literature.

## Acknowledgments

The IRaPPA dataset was a courtesy of the method’s authors Iain H. Moal and Juan Fernandez-Recio. LC thanks the Supercomputing Laboratory at the King Abdullah University of Science and Technology (KAUST) for technical support and access to the Shaheen facilities. DBB was supported by funding from the AI Initiative at KAUST.

## References

Andreani, J., et al. (2013) InterEvScore: a novel coarse-grained interface scoring function using a multi-body statistical potential coupled to evolution. Bioinformatics, 29, 1742–1749.

Barradas-Bautista, D., et al. (2020) The CASP13-CAPRI targets as case studies to illustrate a novel scoring pipeline integrating CONSRANK with clustering and interface analyses. BMC Bioinformatics, 21, 262.

Blum, A.L. and Langley, P. (1997) Selection of relevant features and examples in machine learning. Artificial Intelligence, 97, 245–271.

Cao, Y. and Shen, Y. (2020) Bayesian Active Learning for Optimization and Uncertainty Quantification in Protein Docking. Journal of Chemical Theory and Computation, 16, 5334–5347.

Cao, Y. and Shen, Y. (2020) Energy-based graph convolutional networks for scoring protein docking models. Proteins, 88, 1091–1099.

Chen, R., et al. (2003) ZDOCK: an initial-stage protein-docking algorithm. Proteins, 52, 80–87.

Cheng, F., et al. (2021) Comprehensive characterization of protein-protein interactions perturbed by disease mutations. Nat Genet, 53, 342–353.

Cheng, T.M., et al. (2007) pyDock: electrostatics and desolvation for effective scoring of rigid-body protein-protein docking. Proteins, 68, 503–515.

Chermak, E., et al. (2016) Introducing a Clustering Step in a Consensus Approach for the Scoring of Protein-Protein Docking Models. PLoS One, 11, e0166460.

Chermak, E., et al. (2015) CONSRANK: a server for the analysis, comparison and ranking of docking models based on inter-residue contacts. Bioinformatics, 31, 1481–1483.

de Vries, S.J., et al. (2007) HADDOCK versus HADDOCK: new features and performance of HADDOCK2.0 on the CAPRI targets. Proteins, 69, 726–733.

Dominguez, C., et al. (2003) HADDOCK: A Protein−Protein Docking Approach Based on Biochemical or Biophysical Information. Journal of the American Chemical Society, 125, 1731–1737.

Gabb, H.A., et al. (1997) Modelling protein docking using shape complementarity, electrostatics and biochemical information11Edited by J. Thornton. Journal of Molecular Biology, 272, 106–120.

Garcia-Garcia, J., et al. (2010) Biana: a software framework for compiling biological interactions and analyzing networks. BMC Bioinformatics, 11, 56.

Geng, C., et al. (2020) iScore: a novel graph kernel-based function for scoring protein–protein docking models. Bioinformatics, 36, 112–121.

Grosdidier, S. and Fernández-Recio, J. (2008) Identification of hot-spot residues in protein-protein interactions by computational docking. BMC Bioinformatics, 9, 447.

Harmalkar, A. and Gray, J.J. (2021) Advances to tackle backbone flexibility in protein docking. Curr Opin Struct Biol, 67, 178–186.

Huang, S.Y. (2014) Search strategies and evaluation in protein-protein docking: principles, advances and challenges. Drug Discov Today, 19, 1081–1096.

Hunter, J.D. (2007) Matplotlib: A 2D Graphics Environment. Computing in Science & Engineering, 9, 90–95.

Hwang, H., et al. (2010) Protein-protein docking benchmark version 4.0. Proteins, 78, 3111–3114.

John, G.H., et al. (1994) Irrelevant Features and the Subset Selection Problem. In: Cohen, W.W. and Hirsh, H., editors, Machine Learning Proceedings 1994. San Francisco (CA): Morgan Kaufmann; p. 121–129.

Kastritis, P.L., et al. (2014) Proteins feel more than they see: fine-tuning of binding affinity by properties of the non-interacting surface. J Mol Biol, 426, 2632–2652.

Kudo, M. and Sklansky, J. (2000) Comparison of algorithms that select features for pattern classifiers. Pattern recognition, 33, 25–41.

Lensink, M.F., et al. (2019) Blind prediction of homo- and hetero-protein complexes: The CASP13-CAPRI experiment. Proteins, 87, 1200–1221.

Lensink, M.F., et al. (2018) The challenge of modeling protein assemblies: the CASP12-CAPRI experiment. Proteins, 86 Suppl 1, 257–273.

Lensink, M.F., et al. (2016) Prediction of homoprotein and heteroprotein complexes by protein docking and template-based modeling: A CASP-CAPRI experiment. Proteins, 84 Suppl 1, 323–348.

Liu, S. and Vakser, I.A. (2011) DECK: Distance and environment-dependent, coarse-grained, knowledge-based potentials for protein-protein docking. BMC Bioinformatics, 12, 280.

Lu, H., et al. (2003) Development of unified statistical potentials describing protein-protein interactions. Biophys J, 84, 1895–1901.

Lu, H., et al. (2020) Recent advances in the development of protein–protein interactions modulators: mechanisms and clinical trials. Signal Transduction and Targeted Therapy, 5, 213.

Marcano-Cedeno, A., et al. Feature selection using Sequential Forward Selection and classification applying Artificial Metaplasticity Neural Network. In, IECON 2010 - 36th Annual Conference of IEEE Industrial Electronics. IEEE; 2010. p. 2845–2850.

Mendez, R., et al. (2003) Assessment of blind predictions of protein-protein interactions: current status of docking methods. Proteins, 52, 51–67.

Mitternacht, S. (2016) FreeSASA: An open source C library for solvent accessible surface area calculations. F1000Res, 5, 189.

Moal, I.H., et al. (2017) IRaPPA: information retrieval based integration of biophysical models for protein assembly selection. Bioinformatics, 33, 1806–1813.

Moal, I.H., et al. (2015) Inferring the microscopic surface energy of protein-protein interfaces from mutation data. Proteins, 83, 640–650.

Moal, I.H., et al. (2015) CCharPPI web server: computational characterization of protein-protein interactions from structure. Bioinformatics, 31, 123–125.

Moal, I.H., et al. (2013) Scoring functions for protein-protein interactions. Curr Opin Struct Biol, 23, 862–867.

Moal, I.H., et al. (2013) The scoring of poses in protein-protein docking: current capabilities and future directions. BMC Bioinformatics, 14, 286.

Mosca, R., et al. (2013) Interactome3D: adding structural details to protein networks. Nat Methods, 10, 47–53.

Nadalin, F. and Carbone, A. (2018) Protein-protein interaction specificity is captured by contact preferences and interface composition. Bioinformatics, 34, 459–468.

Oliva, R., et al. (2015) Analysis and Ranking of Protein-Protein Docking Models Using Inter-Residue Contacts and Inter-Molecular Contact Maps. Molecules, 20, 12045–12060.

Oliva, R., et al. (2013) Ranking multiple docking solutions based on the conservation of inter-residue contacts. Proteins, 81, 1571–1584.

Pedregosa, F., et al. (2011) Scikit-learn: Machine Learning in Python. Journal of Machine Learning Research, 12, 2825–2830.

Pokarowski, P., et al. (2005) Inferring ideal amino acid interaction forms from statistical protein contact potentials. Proteins, 59, 49–57.

Pons, C., et al. (2011) Scoring by intermolecular pairwise propensities of exposed residues (SIPPER): a new efficient potential for protein-protein docking. J Chem Inf Model, 51, 370–377.

Rodrigues, J.P., et al. (2012) Clustering biomolecular complexes by residue contacts similarity. Proteins, 80, 1810–1817.

Sahni, N., et al. (2015) Widespread macromolecular interaction perturbations in human genetic disorders. Cell, 161, 647–660.

Schenk, J., et al. Selecting features in on-line handwritten whiteboard note recognition: SFS or SFFS? In.: IEEE; 2009. p. 1251–1254.

Vangone, A. and Bonvin, A.M. (2015) Contacts-based prediction of binding affinity in protein-protein complexes. Elife, 4, e07454.

Vangone, A., et al. (2013) Using a consensus approach based on the conservation of inter-residue contacts to rank CAPRI models. Proteins, 81, 2210–2220.

Vangone, A., et al. (2012) CONS-COCOMAPS: a novel tool to measure and visualize the conservation of inter-residue contacts in multiple docking solutions. BMC Bioinformatics, 13 Suppl 4, S19.

Vangone, A., et al. (2017) Prediction of Biomolecular Complexes. In: J. Rigden, D., editor, From Protein Structure to Function with Bioinformatics. Dordrecht: Springer Netherlands; p. 265–292.

Vangone, A., et al. (2011) COCOMAPS: a web application to analyze and visualize contacts at the interface of biomolecular complexes. Bioinformatics, 27, 2915–2916.

Varoquaux, G., et al. (2015) Scikit-learn.

Vreven, T., et al. (2011) Integrating atom-based and residue-based scoring functions for protein-protein docking. Protein Sci, 20, 1576–1586.

Vreven, T., et al. (2015) Updates to the Integrated Protein-Protein Interaction Benchmarks: Docking Benchmark Version 5 and Affinity Benchmark Version 2. J Mol Biol, 427, 3031–3041.

Wang, X., et al. (2020) Protein docking model evaluation by 3D deep convolutional neural networks. Bioinformatics, 36, 2113–2118.

Waskom, M., et al. Mwaskom/Seaborn: V0.9.0 (July 2018). In.: Zenodo; 2018.

Zhou, H. and Skolnick, J. (2011) GOAP: a generalized orientation-dependent, all-atom statistical potential for protein structure prediction. Biophys J, 101, 2043–2052.

