## Supplementary material for "A Random Forest Classifier for Protein-Protein Docking Models": Barradas-Bautista_CoDES_SI.pdf

**Table S1**. CAPRI criteria for the models assessment, adapted from Lensink et al. (2016) *Proteins*, **84 Suppl 1**, 323-48 (Lensink, et al., 2016).

| **f(nat)** | | **L-rms (Å)** | | **I-rms (Å)** | **Assessment** |
|---|---|---|---|---|---|
| ≥ 0.5 | AND | ≤ 1.0 | OR | ≤ 1.0 | High |
| ≥ 0.3 | AND | < 1.0-5.0] | OR | < 1.0-2.0] | Medium |
| ≥ 0.1 | AND | < 5.0-10.0] | OR | < 2.0-4.0] | Acceptable |
| < 0.1 | AND | > 10.0 | OR | > 4.0 | Incorrect |

**Table S2.** Complete list of features used for the initial classifiers training. For each feature, the public source, relative reference and a short description are provided.

| **Feature** | **Source** | **Reference** | **Description** |
|---|---|---|---|
| ACE_COUL | CCharPPI | J. Phys. Chem. 100:1578 (1996) as calculated using CHARMM. | The change in Coulombic energy using the ACE model |
| ACE_HYDR | CCharPPI | J. Phys. Chem. 100:1578 (1996) as calculated using CHARMM. | The change in hydrophobic energy using the ACE model |
| ACE_INTE | CCharPPI | J. Phys. Chem. 100:1578 (1996) as calculated using CHARMM. | The change in interaction energy using the ACE model |
| ACE_SCRE | CCharPPI | J. Phys. Chem. 100:1578 (1996) as calculated using CHARMM. | The change in screening energy using the ACE model |
| ACE_SELF | CCharPPI | J. Phys. Chem. 100:1578 (1996) as calculated using CHARMM. | The change in self energy using the ACE model |
| ACE_SOLV | CCharPPI | J. Phys. Chem. 100:1578 (1996) as calculated using CHARMM. | The change in solvation energy (sum of ACE_SCRE and ACE_SELF) using the ACE model |
| ALIPH | CCharPPI | Proteins 69(1):139 (2007). | The aliphatic potential integrated in FireDock |
| AP_ACE | CCharPPI | Proteins 69(1):139 (2007). | Contact Energies (J. Mol. Biol. 267(3):707 (1997)) as integrated in FireDock |
| AP_calRW | CCharPPI | PLoS One. 2010 5(10):e15386. | The calRW distance-dependent atomic potential |
| AP_calRWp | CCharPPI | PLoS One. 2010 5(10):e15386. | The calRWplus orientation-dependent atomic potential |

| | | | |
|---|---|---|---|
| AP_DARS | CCharPPI | Biophys J. 2008 95(9):4217-27 | The DARS potential |
| AP_DCOMPLEX | CCharPPI | Proteins 56:93 (2004). | The DComplex potential |
| AP_dDFIRE | CCharPPI | Proteins 72:793 (2008). | Interaction energy calculated using the dDFIRE potential |
| AP_DDG_U | CCharPPI | J. Chem. Theory Comput. 9(8):3715-3727 (2013). | The unweighted atomic potential derived from mutation data |
| AP_DDG_W | CCharPPI | J. Chem. Theory Comput. 9(8):3715-3727 (2013). | The weighted atomic potential derived from mutation data |
| AP_DFIRE2 | CCharPPI | Protein Sci. 17:1212 (2008). | Interaction energy calculated using the DFIRE2 potential |
| AP_DOPE | CCharPPI | Protein Sci. 15(11):2507 (2006). | The DOPE statistical potential |
| AP_DOPE_HR | CCharPPI | Protein Sci. 15(11):2507 (2006). | The high resolution DOPE statistical potential |
| AP_GEOMETRIC | CCharPPI | http://gila.bioengr.uic.edu/resources/geometric.html | The unpublished geometric potential of Li X. and Liang J. |
| AP_GOAP_ALL | CCharPPI | Biophys J. 2011 101(8): 2043-2052. | The total GOAP energy |
| AP_GOAP_DF | CCharPPI | Biophys J. 2011 101(8): 2043-2052. | The DFIRE term in the GOAP energy |
| AP_GOAP_G | CCharPPI | Biophys J. 2011 101(8): 2043-2052. | The GOAP_ag term in the GOAP energy |
| AP_MPS | CCharPPI | Biophys J. 2008 95(9):4217-27 | The MPS potential |
| AP_OPUS_PSP | CCharPPI | J. Mol. Biol. 376:288 (2008). | The OPUS-PSP potential |
| AP_PISA | CCharPPI | Proteins 81(4):592 (2013). | The PISA score |
| AP_T1 | CCharPPI | BMC Struct biol 10:40 (2010). | The first atomic two-step potential |
| AP_T2 | CCharPPI | BMC Struct biol 10:40 (2010). | The second atomic two-step potential |
| AP_URS | CCharPPI | Biophys J. 2008 95(9):4217-27 | The URS potential |
| AP_W1 | CCharPPI | Proteins 2007 69(3):511-20. | A reimplementation of the potential |
| CP_BFKV | CCharPPI | Proteins 59(1):49 (2005) and BMC Bioinformatics 11:92 (2010) | Contact potenital calculated between intermolecular residues. |
| CP_BL | CCharPPI | Proteins 59(1):49 (2005) and BMC Bioinformatics 11:92 (2010) | Contact potenital calculated between intermolecular residues. |
| CP_BT | CCharPPI | Proteins 59(1):49 (2005) and BMC | Contact potenital calculated between intermolecular |

| | | | |
|---|---|---|---|
| | | Bioinformatics 11:92 (2010) | residues. |
| CP_D1 | CCharPPI | BMC Bioinformatics. 2011 12:280. | A reimplementation of the DECK residue level distance-dependent potential |
| CP_DDG_U | CCharPPI | J. Chem. Theory Comput. 9(8):3715-3727 (2013). | The unweighted residue potential derived from mutation data |
| CP_DDG_W | CCharPPI | J. Chem. Theory Comput. 9(8):3715-3727 (2013). | The weighted residue potential derived from mutation data |
| CP_E3D_CB | CCharPPI | Protein Sci. 2011 20(3):529-41. | The E_3D C_beta potential |
| CP_E3D_MIN | CCharPPI | Protein Sci. 2011 20(3):529-41. | The E_3D R_min potential |
| CP_E3DC_CB | CCharPPI | Protein Sci. 2011 20(3):529-41. | The E_3DC C_beta potential |
| CP_E3DC_MIN | CCharPPI | Protein Sci. 2011 20(3):529-41. | The E_3DC R_min potential |
| CP_ELOCAL_CB | CCharPPI | Protein Sci. 2011 20(3):529-41. | The E_local C_beta potential |
| CP_ELOCAL_MIN | CCharPPI | Protein Sci. 2011 20(3):529-41. | The E_local R_min potential |
| CP_EPAIR_CB | CCharPPI | Protein Sci. 2011 20(3):529-41. | The E_pair C_beta potential |
| CP_EPAIR_MIN | CCharPPI | Protein Sci. 2011 20(3):529-41. | The E_pair R_min potential |
| CP_ES3DC_CB | CCharPPI | Protein Sci. 2011 20(3):529-41. | The E_ZS3DC C_beta potential |
| CP_ES3DC_MIN | CCharPPI | Protein Sci. 2011 20(3):529-41. | The E_ZS3DC R_min potential |
| CP_GKS | CCharPPI | Proteins 59(1):49 (2005) and BMC Bioinformatics 11:92 (2010) | Contact potenital calculated between intermolecular residues. |
| CP_HLPL | CCharPPI | Proteins 59(1):49 (2005) and BMC Bioinformatics 11:92 (2010) | Contact potenital calculated between intermolecular residues. |
| CP_MJ1 | CCharPPI | Proteins 59(1):49 (2005) and BMC Bioinformatics 11:92 (2010) | Contact potenital calculated between intermolecular residues. |
| CP_MJ2 | CCharPPI | Proteins 59(1):49 (2005) and BMC Bioinformatics 11:92 (2010) | Contact potenital calculated between intermolecular residues. |
| CP_MJ2h | CCharPPI | Proteins 59(1):49 (2005) and BMC Bioinformatics 11:92 (2010) | Contact potenital calculated between intermolecular residues. |
| CP_MJ3h | CCharPPI | Proteins 59(1):49 (2005) and BMC | Contact potenital calculated between intermolecular |

| | | | |
|---|---|---|---|
| | | Bioinformatics 11:92 (2010) | residues. |
| CP_MJPL | CCharPPI | Proteins 59(1):49 (2005) and BMC Bioinformatics 11:92 (2010) | Contact potenital calculated between intermolecular residues. |
| CP_MS | CCharPPI | Proteins 59(1):49 (2005) and BMC Bioinformatics 11:92 (2010) | Contact potenital calculated between intermolecular residues. |
| CP_MSBM | CCharPPI | Proteins 59(1):49 (2005) and BMC Bioinformatics 11:92 (2010) | Contact potenital calculated between intermolecular residues. |
| CP_Qa | CCharPPI | Proteins 59(1):49 (2005) and BMC Bioinformatics 11:92 (2010) | Contact potenital calculated between intermolecular residues. |
| CP_Qm | CCharPPI | Proteins 59(1):49 (2005) and BMC Bioinformatics 11:92 (2010) | Contact potenital calculated between intermolecular residues. |
| CP_Qp | CCharPPI | Proteins 59(1):49 (2005) and BMC Bioinformatics 11:92 (2010) | Contact potenital calculated between intermolecular residues. |
| CP_RMFCA | CCharPPI | Proteins 65(3):726 (2006). | The C_alpha-C_alpha potential |
| CP_RMFCEN1 | CCharPPI | Proteins 70(3):950 (2006). | The 6bin-HRSC centroid-centroid potential |
| CP_RMFCEN2 | CCharPPI | Proteins 70(3):950 (2006). | The 7bin-HRSC centroid-centroid potential |
| CP_RO | CCharPPI | Proteins 59(1):49 (2005) and BMC Bioinformatics 11:92 (2010) | Contact potenital calculated between intermolecular residues. |
| CP_SJKG | CCharPPI | Proteins 59(1):49 (2005) and BMC Bioinformatics 11:92 (2010) | Contact potenital calculated between intermolecular residues. |
| CP_SKOa | CCharPPI | Proteins 59(1):49 (2005) and BMC Bioinformatics 11:92 (2010) | Contact potenital calculated between intermolecular residues. |
| CP_SKOb | CCharPPI | Proteins 59(1):49 (2005) and BMC Bioinformatics 11:92 (2010) | Contact potenital calculated between intermolecular residues. |
| CP_SKOIP | CCharPPI | Biophys. J. 84(3):1895 (2003). | The residue level interaction contact potential |
| CP_TB | CCharPPI | Proteins 62(4):970 (2006). | The residue level interaction contact potential |
| CP_TD | CCharPPI | Proteins 59(1):49 (2005) and BMC | Contact potenital calculated between intermolecular |

| | | | |
|---|---|---|---|
| | | Bioinformatics 11:92 (2010) | residues. |
| CP_TEl | CCharPPI | Proteins 59(1):49 (2005) and BMC Bioinformatics 11:92 (2010) | Contact potenital calculated between intermolecular residues. |
| CP_TEs | CCharPPI | Proteins 59(1):49 (2005) and BMC Bioinformatics 11:92 (2010) | Contact potenital calculated between intermolecular residues. |
| CP_TS | CCharPPI | Proteins 59(1):49 (2005) and BMC Bioinformatics 11:92 (2010) | Contact potenital calculated between intermolecular residues. |
| CP_TSC | CCharPPI | BMC Struct biol 10:40 (2010). | The residue level interaction two-step potential |
| CP_VD | CCharPPI | Proteins 59(1):49 (2005) and BMC Bioinformatics 11:92 (2010) | Contact potenital calculated between intermolecular residues. |
| CP_Z3DC_CB | CCharPPI | Protein Sci. 2011 20(3):529-41. | The E_3DC Z-score C_beta potential |
| CP_Z3DC_MIN | CCharPPI | Protein Sci. 2011 20(3):529-41. | The E_3DC Z-score R_min potential |
| CP_ZLOCAL_CB | CCharPPI | Protein Sci. 2011 20(3):529-41. | The E_local Z-score C_beta potential |
| CP_ZLOCAL_MIN | CCharPPI | Protein Sci. 2011 20(3):529-41. | The E_local Z-score R_min potential |
| CP_ZPAIR_CB | CCharPPI | Protein Sci. 2011 20(3):529-41. | The E_pair Z-score C_beta potential |
| CP_ZPAIR_MIN | CCharPPI | Protein Sci. 2011 20(3):529-41. | The E_pair Z-score R_min potential |
| CP_ZS3DC_CB | CCharPPI | Protein Sci. 2011 20(3):529-41. | The E_ZS3DC z-score C_beta potential |
| CP_ZS3DC_MIN | CCharPPI | Protein Sci. 2011 20(3):529-41. | The E_ZS3DC z-score R_min potential |
| DDG_V | CCharPPI | Proteins 83(4):640 (2015). | A microscopic surface energy model derived from mutation data |
| DESOLV | CCharPPI | Proteins 68:503 (2007) and Protein 69:852 (2007). | Desolvation energy as calculated using pyDock |
| DOKB | CCharPPI | Biopolymers 101(6):681 (2014). | Energy is calculated by summating the interaction energy between residues which is generated by Boltzmann statistic. DOI:10.1002/bip.22440 |
| ELE | CCharPPI | Proteins 68:503 (2007) and Protein 69:852 (2007). | Total electrostatic energy as calculated using PyDock |
| FIREDOCK | CCharPPI | Proteins 69(1):139 (2007). | The total FireDock energy (default energy function) |

| | | | |
|---|---|---|---|
| FIREDOCK_AB | CCharPPI | Proteins 69(1):139 (2007). | The total FireDock energy (antibody-antigen energy function) |
| FIREDOCK_EI | CCharPPI | Proteins 69(1):139 (2007). | The total FireDock energy (enzyme-inhibitor energy function) |
| HBOND | CCharPPI | Proteins 69(1):139 (2007). | The hydrogen bonding potential integrated in FireDock |
| INSIDE | CCharPPI | Proteins 69(1):139 (2007). | Insideness concavity as integrated in FireDock |
| NSC | CCharPPI | FEBS Lett. 584(6):1163 (2010). | The surface complementarity score |
| ODA | CCharPPI | Int. J. Data Mining Bioinf. 3:55 (2009) and J. Chem. Inf. Mod. 51:370 (2011). | The optimal docking area (ODA) score |
| PI_PI | CCharPPI | Proteins 69(1):139 (2007). | The pi-pi potential integrated in FireDock |
| PROPNSTS | CCharPPI | J. Chem. Inf. Mod. 51:370 (2011). | Amino acid propensity score |
| PYDOCK_TOT | CCharPPI | Proteins 68:503 (2007) and Protein 69:852 (2007). | Total pyDock energy Proteins 68:503 (2007) and Protein 69:852 (2007). |
| ROT_S | CCharPPI | | Change in rotational entropy upon complexation as calculated using CHARMM. |
| SASA | CCharPPI | Proteins 46:24 (2002) | The SASA implicit solvation model as calculated using CHARMM. |
| SIPPER | CCharPPI | J. Chem. Inf. Mod. 51:370 (2011). | The SIPPER potential |
| TRANS_S | CCharPPI | | Change in translational entropy upon complexation as calculated using CHARMM |
| VDW | CCharPPI | Proteins 68:503 (2007) and Proteins 69:852 (2007). | Van der Waals energy as calculated using pyDock |
| ZRANK | CCharPPI | Proteins 67:1078 (2007) | The ZRANK scoring function |
| ZRANK2 | CCharPPI | Proteins 72:270 (2008) | The ZRANK2 scoring function |
| avg_cips_AA | CIPS | Bioinformatics 34:459 (2018) | Sum of CIPS score for Apolar-Apolar contacts over the total number of Apolar-Apolar contacts |
| avg_cips_AAl | CIPS | Bioinformatics 34:459 (2018) | Sum of CIPS score for Apolar-Aliphatic contacts over the total number of Apolar-Aliphatic contacts |
| avg_cips_AAr | CIPS | Bioinformatics 34:459 | Sum of CIPS score for |

| | | | |
|---|---|---|---|
| | | (2018) | Apolar-Aromatic contacts over the total number of Apolar-Aromatic contacts |
| avg_cips_AlAl | CIPS | Bioinformatics 34:459 (2018) | Sum of CIPS score for Aliphatic-Aliphatic contacts over the total number of Aliphatic-Aliphatic contacts |
| avg_cips_AlAr | CIPS | Bioinformatics 34:459 (2018) | Sum of CIPS score for Aliphatic-Aromatic contacts over the total number of Aliphatic-Aromatic contacts |
| avg_cips_ArAr | CIPS | Bioinformatics 34:459 (2018) | Sum of CIPS score for Aromatic-Aromatic contacts over the total number of Aromatic-Aromatic contacts |
| avg_cips_CA | CIPS | Bioinformatics 34:459 (2018) | Sum of CIPS score for Charged-Apolar contacts over the total number of Charged-Apolar contacts |
| avg_cips_CAl | CIPS | Bioinformatics 34:459 (2018) | Sum of CIPS score for Charged-Aliphatic contacts over the total number of Charged-Aliphatic contacts |
| avg_cips_CAr | CIPS | Bioinformatics 34:459 (2018) | Sum of CIPS score for Charged-Aromatic contacts over the total number of Charged-Aromatic contacts |
| avg_cips_CC | CIPS | Bioinformatics 34:459 (2018) | Sum of CIPS score for Charged-Charged contacts over the total number of Charged-Charged contacts |
| avg_cips_CP | CIPS | Bioinformatics 34:459 (2018) | Sum of CIPS score for Charged-Polar contacts over the total number of Charged-Polar contacts |
| avg_cips_PAl | CIPS | Bioinformatics 34:459 (2018) | Sum of CIPS score for Polar-Aliphatic contacts over the total number of Polar-Aliphatic contacts |
| avg_cips_PAr | CIPS | Bioinformatics 34:459 (2018) | Sum of CIPS score for Polar-Aromatic contacts over the total number of Polar-Aromatic contacts |
| avg_cips_PP | CIPS | Bioinformatics 34:459 (2018) | Sum of CIPS score for Polar-Polar contacts over the total number of Polar-Polar contacts |
| cips_AA | CIPS | Bioinformatics 34:459 (2018) | Sum of CIPS score for Apolar-Apolar contacts |
| cips_AAl | CIPS | Bioinformatics 34:459 (2018) | Sum of CIPS score for Apolar-Aliphatic contacts |

| | | | |
|---|---|---|---|
| cips_AAr | CIPS | Bioinformatics 34:459 (2018) | Sum of CIPS score for Apolar-Aromatic contacts |
| cips_AlAl | CIPS | Bioinformatics 34:459 (2018) | Sum of CIPS score for Aliphatic-Aliphatic contacts |
| cips_AlAr | CIPS | Bioinformatics 34:459 (2018) | Sum of CIPS score for Aliphatic-Aromatic contacts |
| cips_ArAr | CIPS | Bioinformatics 34:459 (2018) | Sum of CIPS score for Aromatic-Aromatic contacts |
| cips_CA | CIPS | Bioinformatics 34:459 (2018) | Sum of CIPS score for Charged-Apolar contacts |
| cips_CAl | CIPS | Bioinformatics 34:459 (2018) | Sum of CIPS score for Charged-Alphatic contacts |
| cips_CC | CIPS | Bioinformatics 34:459 (2018) | Sum of CIPS score for Charged-Charged contacts |
| cips_CP | CIPS | Bioinformatics 34:459 (2018) | Sum of CIPS score for Charged-Polar contacts |
| cips_PA | CIPS | Bioinformatics 34:459 (2018) | Sum of CIPS score for Polar-Apolar contacts |
| cips_PAl | CIPS | Bioinformatics 34:459 (2018) | Sum of CIPS score for Polar-Aliphatic contacts |
| cips_PP | CIPS | Bioinformatics 34:459 (2018) | Sum of CIPS score for Polar-Polar contacts |
| avg_cips_PA | CIPS | Bioinformatics 34:459 (2018) | Sum of CIPS score for Polar-Apolar contacts over the total number of Polar-Apolar contacts |
| CONSRANK score | CONSRANK | Bioinformatics 31:1481 (2015)<br>Proteins 81:1571 (2013) | CONSRANK normalized score, reflecting the conservation of the inter-residue contacts of a given model in the decoys set |
| num_of_contacts | CONSRANK | Bioinformatics 31:1481 (2015)<br>Proteins 81:1571 (2013) | Total number of contacts as calculated in ConsRank |
| AA | COCOMAPS | Bioinformatics 27:2915 (2011) | Apolar-Apolar contact count at 5 Å distance |
| AA_sqrt | COCOMAPS | Bioinformatics 27:2915 (2011) | Square Root of Apolar-Apolar contact count at 5 Å distance |
| AAl | COCOMAPS | Bioinformatics 27:2915 (2011) | Apolar-Aliphatic contact count at 5 Å distance |
| AAl_sqrt | COCOMAPS | Bioinformatics 27:2915 (2011) | Square Root of Apolar-Aliphatic contact count at 5 Å distance |
| AAr | COCOMAPS | Bioinformatics 27:2915 (2011) | Apolar-Aromatic contact count at 5 Å distance |
| AAr_sqrt | COCOMAPS | Bioinformatics 27:2915 (2011) | Square Root of Apolar-Aromatic contact count at 5 Å distance |
| AlAl | COCOMAPS | Bioinformatics 27:2915 | Aliphatic-Aliphatic contact |

| | | | |
|---|---|---|---|
| | | (2011) | count at 5 Å distance |
| AlAl_sqrt | COCOMAPS | Bioinformatics 27:2915 (2011) | Square Root of Aliphatic-Aliphatic contact count at 5 Å distance |
| AlAr | COCOMAPS | Bioinformatics 27:2915 (2011) | Aliphatic-Aromatic contact count at 5 Å distance |
| AlAr_sqrt | COCOMAPS | Bioinformatics 27:2915 (2011) | Square Root of Aliphatic-Aromatic contact count at 5 Å distance |
| ArAr | COCOMAPS | Bioinformatics 27:2915 (2011) | Aromatic-Aromatic contact count at 5 Å distance |
| ArAr_sqrt | COCOMAPS | Bioinformatics 27:2915 (2011) | Square Root of Aromatic-Aromatic contact count at 5 Å distance |
| CA | COCOMAPS | Bioinformatics 27:2915 (2011) | Charged-Apolar contact count at 5 Å distance |
| CA_sqrt | COCOMAPS | Bioinformatics 27:2915 (2011) | Square Root of Charged-Apolar contact count at 5 Å distance |
| CAl | COCOMAPS | Bioinformatics 27:2915 (2011) | Charged-Aliphatic contact count at 5 Å distance |
| CAl_sqrt | COCOMAPS | Bioinformatics 27:2915 (2011) | Square Root of Charged-Aliphatic contact count at 5 Å distance |
| CAr | COCOMAPS | Bioinformatics 27:2915 (2011) | Charged-Aromatic contact count at 5 Å distance |
| CAr_sqrt | COCOMAPS | Bioinformatics 27:2915 (2011) | Square Root of Charged-Aromatic contact count at 5 Å distance |
| CC | COCOMAPS | Bioinformatics 27:2915 (2011) | Charged-Charged contact count at 5 Å distance |
| CC_sqrt | COCOMAPS | Bioinformatics 27:2915 (2011) | Square Root of Charged-Charged contact count at 5 Å distance |
| CP | COCOMAPS | Bioinformatics 27:2915 (2011) | Charged-Polar contact count at 5 Å distance |
| CP_sqrt | COCOMAPS | Bioinformatics 27:2915 (2011) | Square Root of Charged-Polar contact count at 5 Å distance |
| PA | COCOMAPS | Bioinformatics 27:2915 (2011) | Polar-Apolar contact count at 5 Å distance |
| PA_sqrt | COCOMAPS | Bioinformatics 27:2915 (2011) | Square Root of Polar-Apolar contact count at 5 Å distance |
| PAl | COCOMAPS | Bioinformatics 27:2915 (2011) | Polar-Aliphatic contact count at 5 Å distance |
| PAl_sqrt | COCOMAPS | Bioinformatics 27:2915 (2011) | Square Root of Polar-Aliphatic contact count at 5 Å distance |
| PAr | COCOMAPS | Bioinformatics 27:2915 (2011) | Polar-Aromatic contact count at 5 Å distance |
| PAr_sqrt | COCOMAPS | Bioinformatics 27:2915 (2011) | Square Root of Polar-Aromatic contact count at 5 Å distance |

| | | | |
|---|---|---|---|
| PP | COCOMAPS | Bioinformatics 27:2915 (2011) | Polar-Polar contact count at 5 Å distance |
| PP_sqrt | COCOMAPS | Bioinformatics 27:2915 (2011) | Square Root of Polar-Polar contact count at 5 Å distance |
| BSA | Freesasa | F1000Research 5:189 (2016) | Total Buried Surface Area from FreeSASA |
| BSA_Apolar | Freesasa | F1000Research 5:189 (2016) | Polar Buried Surface Area from FreeSASA |
| BSA_Polar | Freesasa | F1000Research 5:189 (2016) | Apolar Buried Surface Area from FreeSASA |
| Nis_Apolar | Prodigy | eLife 4: 291 (2015) J Mol Biol 426: 2632 (2014) | Percentage of charged non-interacting surface (NIS) on residues from Binding affinity predictor based on Intermolecular Contacts (ICs). |
| Nis_Polar | Prodigy | eLife 4: 291 (2015) J Mol Biol 426: 2632 (2014) | Percentage of apolar non-interacting surface (NIS) on residues from Binding affinity predictor based on Intermolecular Contacts (ICs). |

**Table S3.** Global parameters: Accuracy (Acc), Recall (R), Precision (P), F1_score (F1), Matthews' correlation coefficient (MCC) for all the discussed classifiers. These values are also reported in the form of radar plots in the main text.

| Classifier | Acc | R_inc | R_cor | P_inc | P_corr | F1_inc | F1_cor | MCC | Validation set | Figs |
|---|---|---|---|---|---|---|---|---|---|---|
| RF | 0.8321 | 0.8669 | 0.7973 | 0.8105 | 0.8569 | 0.8377 | 0.8260 | 0.6658 | Bal-BM5-up | 2,3 |
| PRC | 0.8184 | 0.8274 | 0.8094 | 0.8128 | 0.8242 | 0.8200 | 0.8168 | 0.6369 | Bal-BM5-up | 2 |
| SVM | 0.7170 | 0.6863 | 0.7477 | 0.7312 | 0.7045 | 0.7080 | 0.7254 | 0.4348 | Bal-BM5-up | 2 |
| RF | 0.9164 | 0.9232 | 0.7985 | 0.9874 | 0.3777 | 0.9543 | 0.5128 | 0.5133 | 3K-BM5-up | 2,3,5 |
| PRC | 0.8683 | 0.8726 | 0.7957 | 0.9865 | 0.2670 | 0.9261 | 0.3998 | 0.4116 | 3K-BM5-up | 2 |
| SVM | 0.7085 | 0.7072 | 0.7299 | 0.9782 | 0.1270 | 0.8209 | 0.2163 | 0.2144 | 3K-BM5-up | 2 |
| RF | 0.8222 | 0.9246 | 0.7199 | 0.768 | 0.9058 | 0.8388 | 0.8017 | 0.659 | Bal-10CV-BM5 | 3 |
| RF | 0.9511 | 0.9624 | 0.7341 | 0.9859 | 0.5043 | 0.974 | 0.5973 | 0.584 | 3K-BM5-up | 3 |
| CoDES | 0.9183 | 0.9197 | 0.8989 | 0.9916 | 0.4618 | 0.9543 | 0.6101 | 0.6093 | 3K-BM5-up | 5 |
| CoDES | 0.7073 | 0.6712 | 0.9852 | 0.9971 | 0.2803 | 0.8023 | 0.4364 | 0.4267 | Score_set | 5 |
| CoDES | 0.6937 | 0.6216 | 0.9948 | 0.9980 | 0.3864 | 0.7661 | 0.5566 | 0.4867 | Score_set (>2.5%) | 5 |

**Table S4.** Features with the highest importance are reported for the RF classifier trained on the balanced sets with a BM4/BM5-update and 10-fold cross-validation approach. Features are sorted based on values of the second column. The mean importance value and associated standard deviation is reported in the last column. The 16 features selected because having an importance above 0.01 for both the classifiers were selected – they are reported in bold. Values below the importance threshold of 0.01 and the corresponding features are reported in italics.

| Feature | Bal-BM4/5-up | Bal-10CV-BM5 |
|---|---|---|
| **CONSRANK_val** | 0.2056 | 0.1978+/-0.0074 |
| **CP_HLPL** | 0.0225 | 0.0256+/-0.0018 |
| **CP_SKOIP** | 0.0172 | 0.0233+/-0.0039 |
| **PYDOCK_TOT** | 0.0223 | 0.0232+/-0.0022 |
| **CP_MJ3h** | 0.0224 | 0.0229+/-0.0031 |
| **DDG_V** | 0.0225 | 0.0226+/-0.0021 |
| **ELE** | 0.0177 | 0.0192+/-0.0013 |
| **SIPPER** | 0.0149 | 0.0171+/-0.0029 |
| **AP_GOAP_DF** | 0.0275 | 0.0168+/-0.0027 |
| **CP_D1** | 0.024 | 0.0151+/-0.0025 |
| *CP_TB* | *0.0093* | 0.0141+/-0.0024 |
| **AP_PISA** | 0.0118 | 0.0138+/-0.0025 |
| **CP_TD** | 0.0247 | 0.0136+/-0.0013 |
| **CP_RMFCA** | 0.0101 | 0.0129+/-0.0015 |
| **AP_dDFIRE** | 0.012 | 0.0127+/-0.0026 |
| **CP_TSC** | 0.0101 | 0.0115+/-0.0014 |
| **AP_DFIRE2** | 0.0129 | 0.0113+/-0.0018 |
| *AP_DARS* | *0.0075* | 0.0105+/-0.0018 |
| *CP_BT* | *0.0083* | 0.0104+/-0.0029 |
| *CP_MJ2h* | 0.017 | *0.0089+/-0.0023* |
